## Supplementary Figures for "Induction of selective cell death in HIV-1-infected cells by DDX3 inhibitors leads to depletion of the inducible reservoir"

**SUPPLEMENTARY INFORMATION**

Supplementary fig. 1

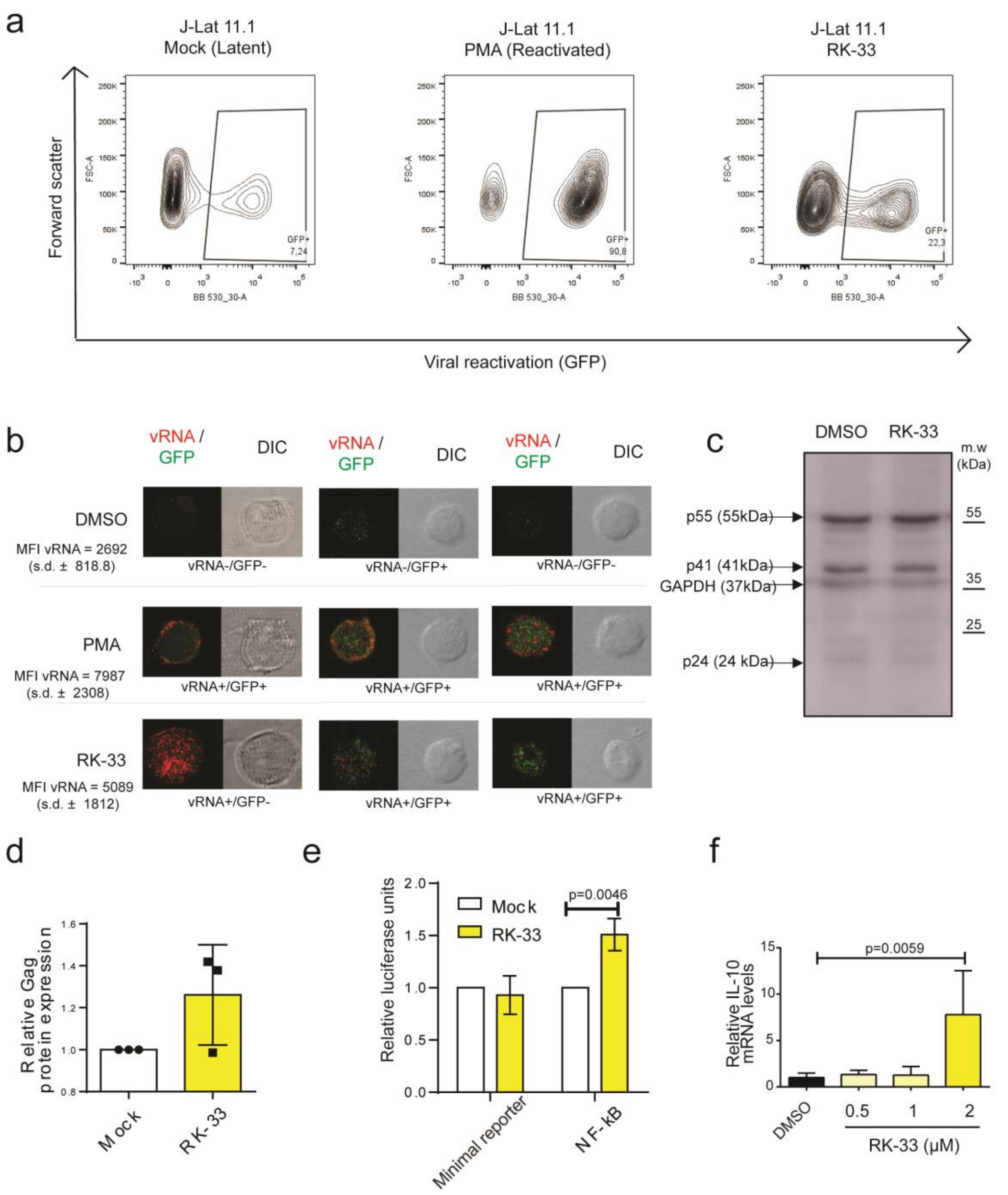


**Fig. 1. Effects of RK-33 on J-Lat 11.1 cells**

**a** Contour plots depicting % GFP expression of Mock, PMA and 2µM RK-33 treated J-Lat 11.1 cells. **b** Three representative images from each condition of J-Lat cells as treated in (a), seeded as unsorted whole populations onto a coverslip and observed by confocal microscopy. vRNA is depicted in red, GFP in green and the differential interference contrast (DIC) in grey. Mean Fluorescence intensity (MFI) of the vRNA channel from three independent experiments are also quantitated and shown. **c** Representative Western blots depicting viral proteins Gag (p55), p41 and p24, and GAPDH in mock and RK-33 treated J-Lat 11.1 cells. **d** Quantification of the densitometric analysis of Gag protein expression normalised to loading control GAPDH. Error bars represent mean ± SD from three independent experiments (Unpaired, two-tailed *t*-test). **e** Relative luciferase expression from cells transfected with a minimal reporter plasmid or those containing NF-κB target sites normalised to mock-treated condition. Error bars represent mean ± SD from three independent experiments (Unpaired, two-tailed *t*-test). **f** The relative mRNA expression levels of IL-10 in J-Lat 11.1 cells treated with increasing concentrations of RK-33 was quantified by RT-qPCR. Error bars represent mean ± SD from three independent experiments (Unpaired, two-tailed *t*-test).

Supplementary fig. 2

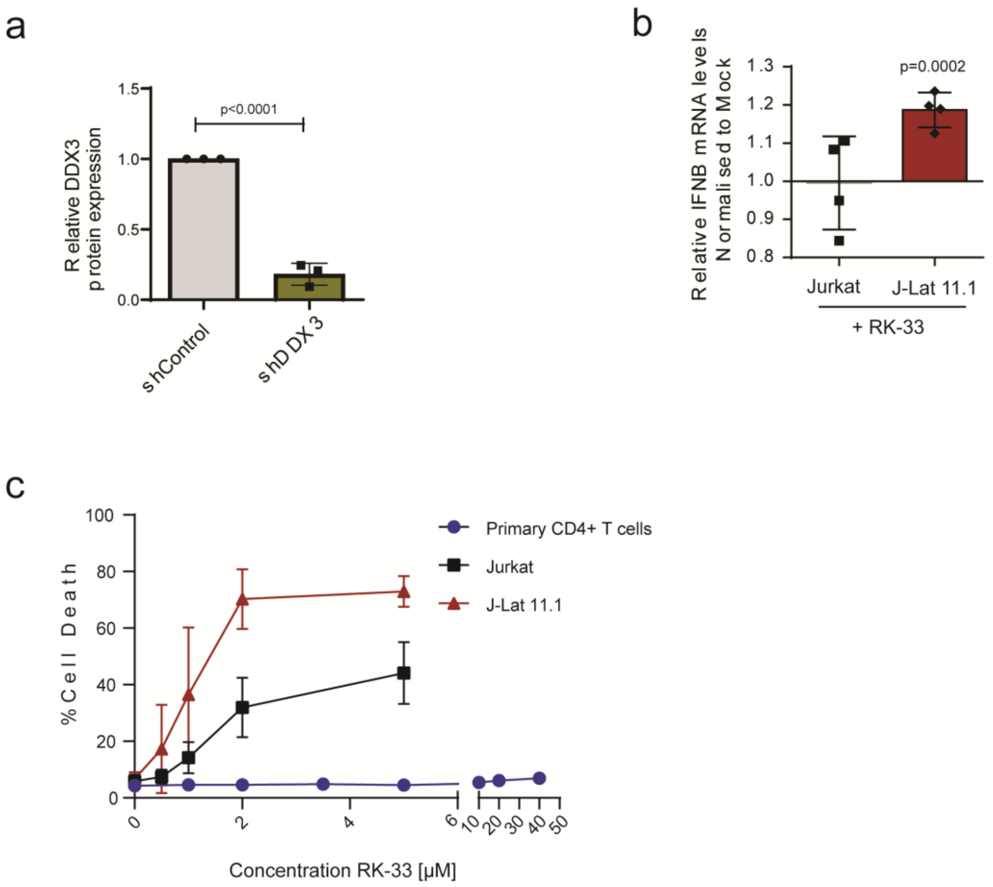


**Fig. 2. Effects of DDX3 inhibition and vRNA expression in J-Lat 11.1 cells**

**a** Quantification of the densitometric analysis of DDX3 expression from Fig. 2b. normalised to loading control GAPDH. Error bars represent mean ± SD from three independent experiments (Unpaired, two-tailed *t*-test). **b** Jurkat and J-Lat 11.1 cells were treated with 2uM RK-33 and relative IFNβ mRNA expression levels were quantified by RT-qPCR and normalised to the mock-treated cells. Error bars represent mean ± SD from three independent experiments (Unpaired, two-tailed *t*-test). **c** % Cell death was measured in Jurkat, J-Lat and primary CD4+ T cells treated with increasing concentrations of RK-33 by flow cytometry. Error bars represent mean ± SD from three independent experiments for cell lines and from 3 independent donors for primary cells.

Supplementary fig. 3

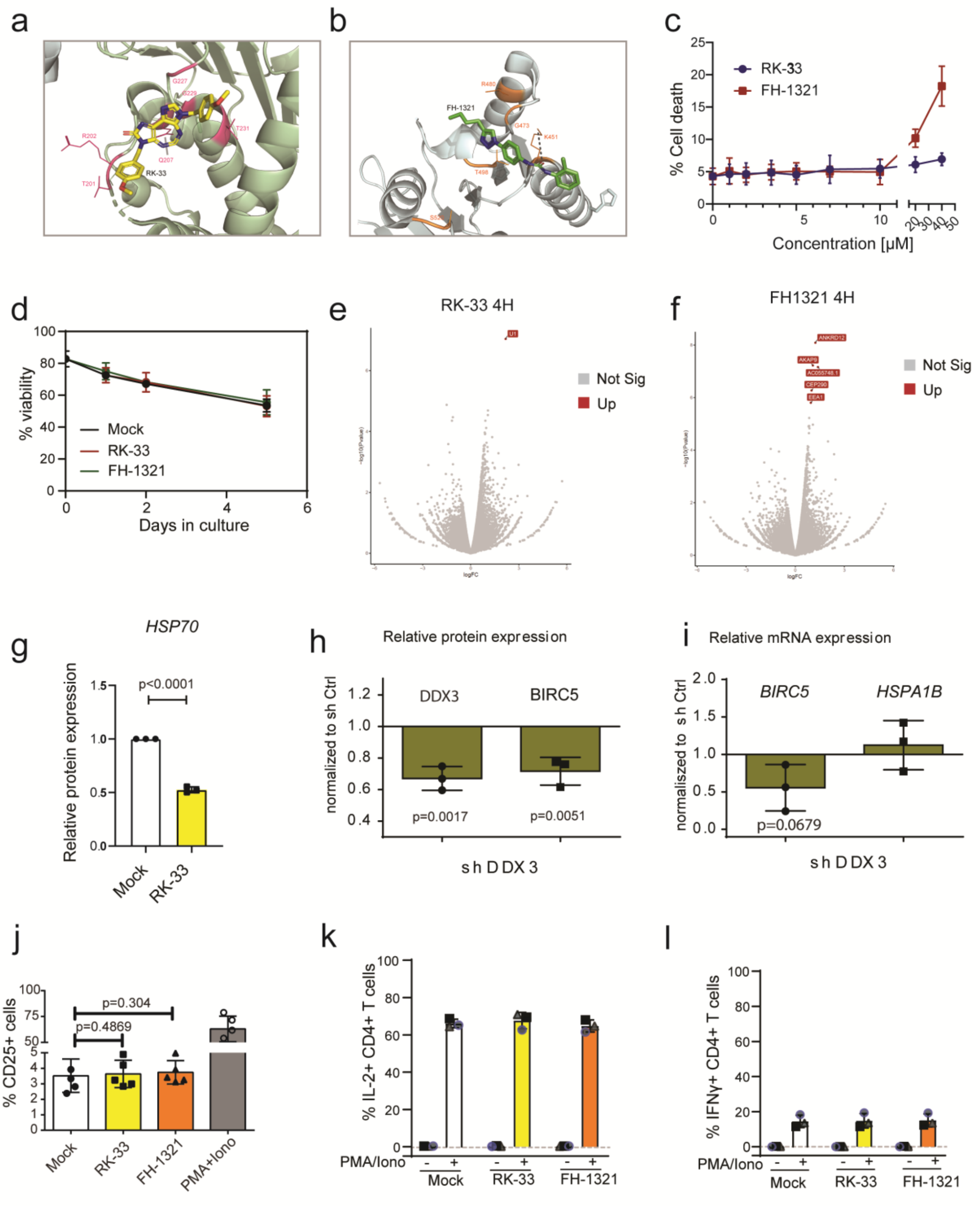


**Fig. 3. Effects of DDX3 inhibitors on primary CD4+ T cells**

**a** Close up of ATP-binding pocket on the surface of DDX3 with bound RK-33 as predicted by AutoDock Vina. DDX3 domain 1 is shown as pale green cartoon, domain 2 is shown as pale blue cartoon and surface color code is pink for ATP-binding residues and orange for RNA-binding residues. Color code for RK33 is yellow for carbon, red for oxygen, dark blue for nitrogen. **b** Close up of RNA-binding pocket on the surface of DDX3 with bound FH-1321. Color code for FH-1321 is green for carbon, red for oxygen, dark blue for nitrogen. The predicted hydrogen bond between FH-1321 and Lysine 541 from DDX3 RNA-binding pocket is indicated with a dashed line. **c** % Cell death of CD4+ T cells treated with increasing concentrations of either RK-33 or FH-1321. Error bars represent mean ± SD from freshly isolated CD4+ T cells from three healthy donors. **d** % live cells (cell viability) of CD4+ T cells treated with 2 µM RK-33 or 1 µM FH-1321 for up to 5 days. Error bars represent mean ± SD from CD4+ T cells isolated from freeze-thawed PBMCs from three healthy donors. Volcano plot of differentially expressed genes from CD4+ T cells 4 hours post-treatment with **e** RK-33 and **f** FH-1321. **g** Quantification of the densitometric analysis of HSP70 protein expression from cells as treated in Fig 2h. Error bars represent mean ± SD from three donors. (Unpaired, two-tailed *t*-test). **h** Quantification of the densitometric analysis of DDX3 and BIRC5 protein expression normalised to loading control GAPDH relative to shControl-treated cells. Error bars represent mean ± SD from three donors (Unpaired, two-tailed *t*-test). **i** Relative mRNA expression of BIRC5 and HSPA1B of shDDX3-treated cells normalised to shControl. Error bars represent mean ± SD from three donors (Unpaired, two-tailed *t*-test). **j** CD4+ T cells from healthy donors were treated with DMSO (Mock), RK-33, FH1321 or PMA-Ionomycin and the % of cells expressing the T-cell activation marker CD25 by flow cytometry. Error bars represent mean ± SD from independent experiments from 5 different donors (Paired, two-tailed *t*-test). Percentage of IL-2 **k** and IFNγ **l** producing cells in unstimulated and stimulated primary CD4+ lymphocytes after treatment with RK-33 and FH-1321 (n=3).

Supplementary fig. 4

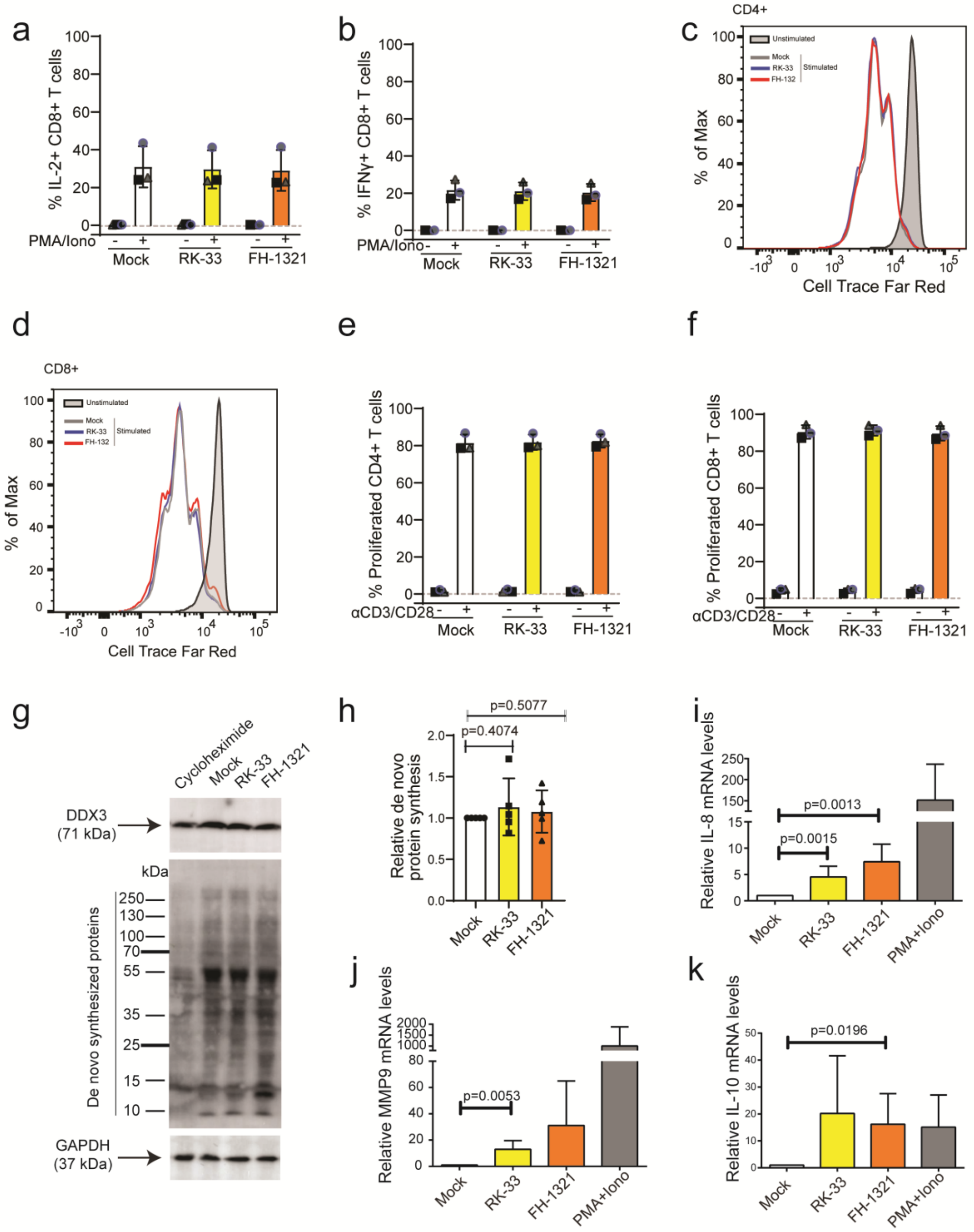


**Fig. 4. DDX3 inhibition has negligible effects on uninfected CD4+ T cells**

Percentage of IL-2 **a** and IFNγ **b** producing CD8+ T cells after treatment with RK-33 and FH-1321. Error bars represent mean ± SD from three donors. Representative histogram overlay of proliferative capacity of unstimulated or aCD3/CD28 stimulated **c** CD4+ T cells or **d** CD8+ T cells in the presence or absence DDX3 inhibitors. Percentage of proliferated **e** CD4+ T cells and **f** CD8+ T cells from 3 healthy donors as described in A. Error bars represent mean ± SD from freshly isolated CD4+ T cells from three donors. **g** Representative Western blots depicting DDX3, GAPDH and de novo synthesized proteins in mock, RK-33, FH-1321 and cycloheximide-treated conditions. **h** Quantification of the densitometric analysis of de novo synthesized proteins normalised to loading control GAPDH. Error bars represent mean ± SD from three donors (Unpaired, two-tailed *t*-test). Relative mRNA expression levels of **i** IL-8, **j** MMP9 and **k** IL-10 from cells treated as in (a) were quantified by RT-qPCR. Error bars represent mean ± SD from independent experiments from at 5 different donors (Unpaired, two-tailed *t*-test).

Supplementary fig. 5

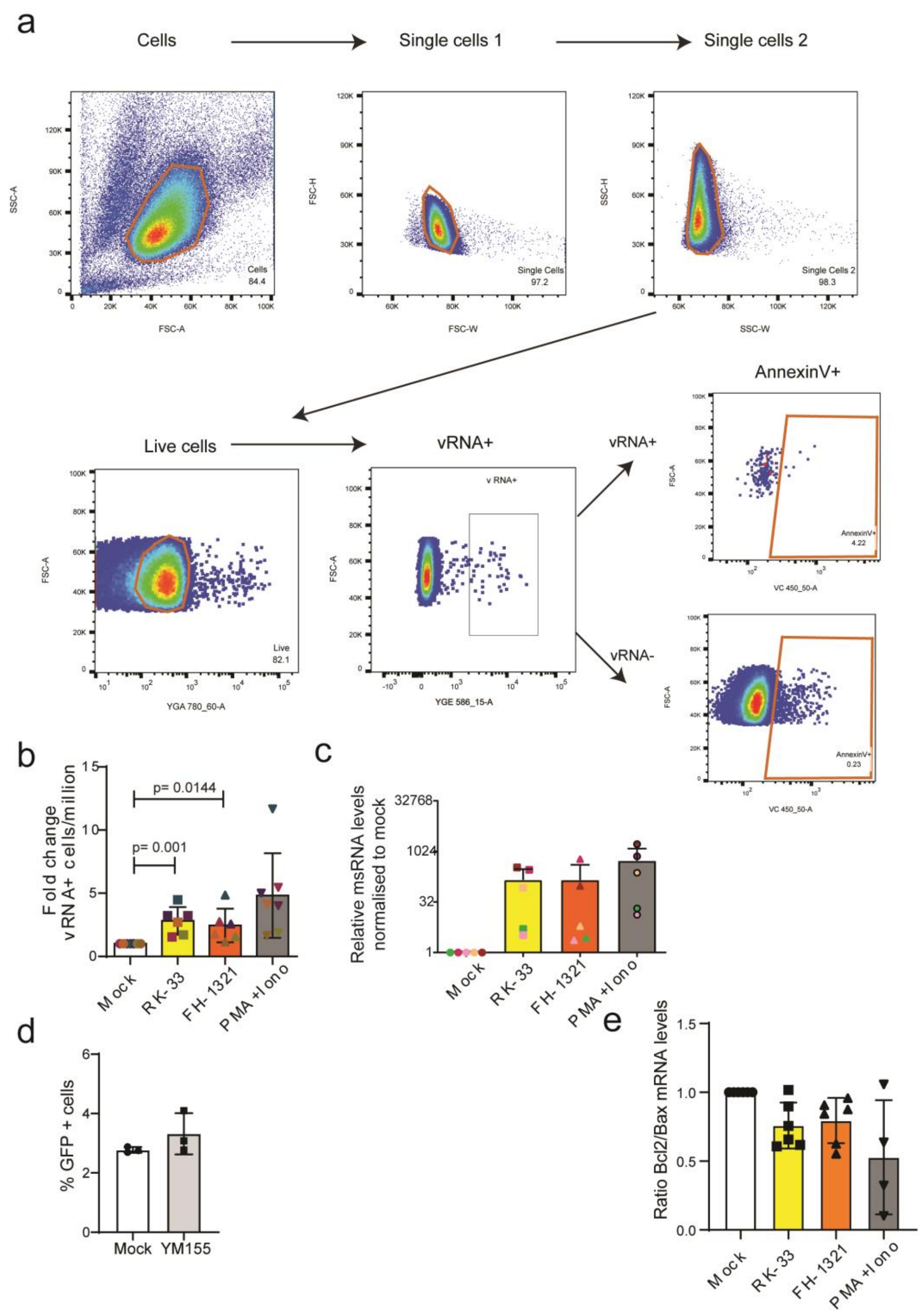


### Fig. 5. DDX3 inhibitors in an *in vitro* model of HIV-1 latency

**a** Gating strategy of FISH-Flow analysis from in vitro-infected CD4+ T cells. Each gate is labelled with a name and % of parent gate. **b** The fold change of number of vRNA-expressing cells per million normalised to the mock-treated condition as depicted in Fig 4c were quantified. Error bars represent mean ± SD from independent experiments from at least 6 different donors, with a minimum of 100,000 events acquired for each condition (Unpaired, two-tailed *t*-test). **c** CD4+ T cells from healthy donors were infected in vitro and treated with DMSO (Mock), RK-33, FH1321 or PMA-Ionomycin. The relative expression of multiply-spliced viral RNA (msRNA) were quantified by RT-qPCR. Error bars represent mean ± SD from independent experiments from 5 different donors. d Treatment of J-Lat 11.1 cells with YM155 does not induce latency reversal as measured by % GFP + cells by flow cytometry from 3 independent experiments. **e** CD4+ T cells from healthy donors were infected in vitro and treated with DMSO (Mock), RK-33, FH1321 or PMA-Ionomycin. The relative expression of ratios of Bcl2 and Bax mRNA expression were quantified by RT-qPCR. Error bars represent mean ± SD from independent experiments from 5 different donors.

Supplementary fig. 6

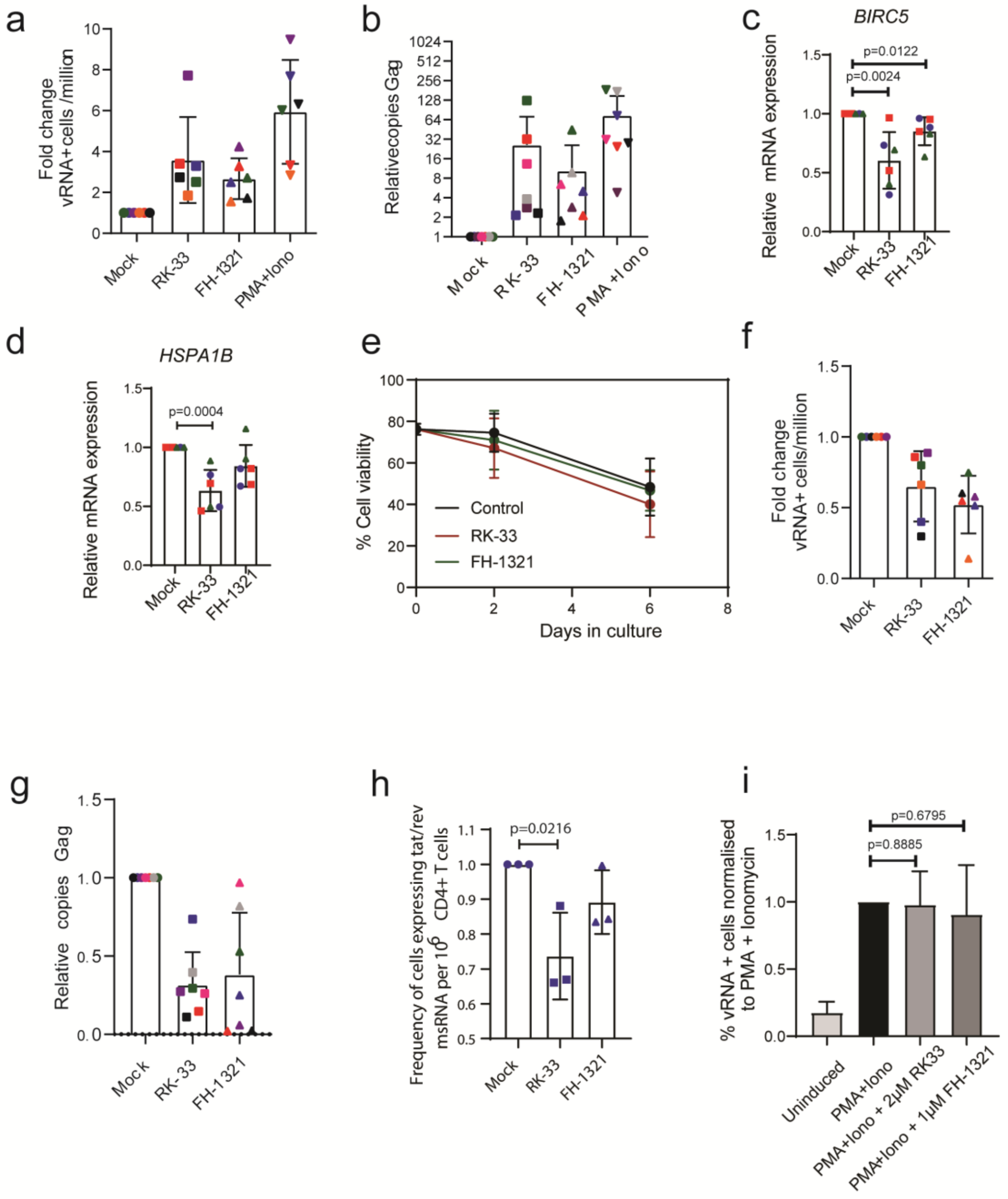


**Fig. 6. DDX3 inhibitors in CD4+ T cells from PLWHIV**

**a** CD4+ T cells isolated from HIV-1-infected donors were treated with DMSO (Unstimulated), PMA/Ionomycin, and the DDX3 inhibitors RK-33 and FH1321. For all graphs, different donors are represented by an individual colour consistently. The fold change of number of vRNA-expressing cells per million normalised to the mock-treated condition as depicted in Fig 5c were quantified. Error bars represent mean ± SD from independent experiments from 6 different donors in duplicate, with a minimum of 100,000 events acquired for each condition. **b** Copies Gag/µg total RNA 18 hours post treatments for each condition was quantified by RT-qPCR and normalised to mock-treated condition. Error bars represent mean ± SD from independent experiments from 7 different donors. Relative mRNA expressions of **c** BIRC5 and **d** HSPA1B was quantified by RT-qPCR. Error bars represent mean ± SD from independent experiments from three donors in duplicate (Unpaired, two-tailed *t*-test). **e** % live cells (cell viability) of CD4+ T cells treated with 2 µM RK-33 or 1 µM FH-1321 for up to 5 days. Error bars represent mean ± SD from CD4+ T cells isolated from freeze-thawed PBMCs from three PLWHIV donors. **f** The fold change of number of vRNA-expressing cells per million normalised to the mock-treated condition as depicted in Fig 6c were quantified. Error bars represent mean ± SD from independent experiments from 6 different donors in duplicate, with a minimum of 100,000 events acquired for each condition (Unpaired, two-tailed *t*-test). **g** Copies Gag/µg total RNA 6 days post treatment post treatments for each condition was quantified by RT-qPCR and normalised to mock-treated condition. Error bars represent mean ± SD from independent experiments from 7 different donors. **h** Fold change of the frequency of MS RNA+ cells as measured by TILDA upon treatment with RK-33 and FH-1321 ,normalised to the Mock-treated cells. Error bars represent mean ± SD of three independent experiments from one donor (Unpaired, two-tailed *t*-test). **i** In vitro HIV-1-infected CD4+ T cells were left uninduced or reactivated with PMA/Ionomycin with or without RK-33 and FH-1321. % vRNA expression normalised to PMA-Ionomycin condition are depicted. Error bars represent mean ± SD from independent experiments from 3 different donors (Unpaired, two-tailed *t*-test).
